# Impacts of climate change on fonio millet: seed germination and suitability modelling of an important indigenous West African crop

**DOI:** 10.1101/2025.03.14.641845

**Authors:** George P. Burton, Hillary Mireku Botey, Paolo Ceci, Caspar Chater, Rafal M. Gutaker, Amy C. Jackson, Philippa Ryan, Charlotte E. Seal, Colin G. N. Turnbull, Maria S. Vorontsova, Efisio Mattana, Tiziana Ulian

## Abstract

Seed germination is highly temperature sensitive. Climate change factors such as increasing temperatures are likely to have a harmful effect on agriculture, particularly after crop sowing. Better utilisation of indigenous, arid-resilient crops like fonio (*Digitaria exilis*) are a commonly proposed solution to improving food security. This study develops knowledge of fonio germination requirements and how these correspond to future climate conditions across West Africa. We use a combined approach; integrating seed germination experiments under a range of temperatures, and niche suitability modelling to investigate how cultivation of fonio will be impacted by climate change. We find that from 37 seed accessions collected across Guinea, Togo, Mali, and Burkina Faso, the ceiling temperature for germination is around 42°C, with an optimum temperature of 30-35°C – also noted from phenotypic observations. Drought trials show successful germination to beyond -1MPa. There is no obvious difference in response by accessions originating from either hotter or cooler climates. By comparing these temperature thresholds with future climate predictions, alongside a suitability modelling approach, we predict an average decline of 10% in the suitable area for fonio cultivation, especially affecting Senegal, Mali, and Burkina Faso. Newly suitable area is predicted to increase in Guinea, Ghana, Cote d’Ivoire, and Nigeria by around 5%. These findings provide valuable insight for developing future dryland agriculture policies and prioritisation of resilient crops.

## Introduction

### Climate change and heat-resilient crops

In past decades there has been a significant increase in both rising global temperatures and unpredictable, destructive weather patterns which affect both the quality and availability of food crops, and overall human health (Sylla et al., 2018; Fatima et al., 2020; Ragatoa et al., 2024). Climate models suggest that an increase of 1-2°C could further destabilise agricultural systems (Sylla et al., 2018), yet overall global temperatures are predicted to surpass this, with increases of up to 2-3°C by 2050 (Fatima et al., 2020), which will be particularly harmful to agriculture in vulnerable arid regions of West Africa. This is concerning for regions that often prioritise major introduced crops such as Asian rice (*Oryza sativa* L.) which requires irrigation and is less resilient to the local high heat and drought than native locally-adapted crops like African rice (*Oryza glaberrima* Steud., Linares, 2002). Other indigenous crops, such as cereal millet fonio (*Digitaria exilis* (Kippist) Stapf.), exhibit high tolerance to heat and drought, making them critical to food systems for future food security in West Africa (Cruz et al., 2016).

Countries close to the Sahel in West Africa have recently experienced record weather conditions, including a heatwave in March-April 2024, with prolonged temperatures above 45°C in Senegal, Mali, and Burkina Faso (World Weather Attribution, April 2024) - this had severe effects on human health causing hospitalisations and deaths. These record temperatures were coupled with extreme rainfall and flooding in the Niger and Lake Chad basins later in Autumn, again causing damage and loss of life (World Weather Attribution, October 2024). Increases in consistent average temperature and frequency of extreme weather events are predicted across the Sahel (Ragatoa et al., 2024).

Seed germination is highly sensitive to ambient temperature, and warming climates are likely to have an adverse effect on both plant germination success and post-germination seedling development. Millets, however, have a higher-than-average heat tolerance to protect against these stresses (Wahid et al., 2007; Tadele et al., 2016), utilising various adaptation strategies to grow in challenging climates (Pirnajmedin et al., 2024; Heckman et al., 2024). Climate change impacts on food security are directly related to seed physiology and vigour: the capacity of a crop species to regulate successful germination and survival under heat and drought stress (Reed et al., 2022). As the climate changes, it will be essential to identify the specific heat and drought limits of germination for indigenous crops, to select the most climate-resilient crops appropriate for each region.

### Seed germination

Seed germination studies have a longstanding history in botanical research. Systematic methods for assessing germination rates became commonplace in the 1980’s, using the proportion of germinated seeds under a set of conditions to predict germination under untested conditions. This involved the concepts of cardinal temperatures: base temperature (Tb), optimum temperature (To), and ceiling temperature (Tc) – to delimit the thresholds within a seed population will germinate under a gradient of different temperatures, to mathematically predict the effects of temperature on seed germination rate (Garcia-Huidobro et al., 1982; Ellis et al., 1986; Gummerson, 1986; Covell et al., 1986). These studies introduced the use of a ‘thermal-time’ model, and later a ‘hydrothermal-time’ model, the latter to predict the effects of water potential and drought on germination. These concepts and models are widely applied (Watt & Bloomberg, 2012; Bradford, 2002; Trudgill et al., 2000), including the use of polyethylene glycol (PEG) to create solutions of different water potential to limit the movement of water into a seed, thereby simulating drought. Though there have been concerns for how PEG may interact with toxicity stress, temperature, and substrates like filter paper (Michel, 1983; Hardegree and Emmerich, 1990), they nonetheless provide an accessible and efficient method of testing the effects of temperature and/or simulated drought (a decrease in water potential) on seed germination through time. More recently, dose-response curves (which rely on a time-to-event model) have been presented as a mathematically accurate method of calculating germination rates which are inherently non-linear, including using non-parametric maximum likelihood estimator (NPMLE) models (Onofri et al., 2018; Catara et al., 2016; Onofri et al., 2022).

### Fonio climate resilience

Fonio is a West African millet known to tolerate extreme drought and high temperatures (Tadele et al., 2016; Cruz et al., 2016), as well as grow in unfertilised, un-irrigated soils, and has an important cultural role in religious and social ceremonies (Burton et al., 2023). It is therefore an important crop in supporting food security of rural communities in West Africa (Cruz et al., 2016). Despite its important role in sustainable, subsistence agriculture and potential for improving livelihoods in rural communities, fonio has been recognised as a Neglected and Underutilised Species (NUS, Ulian et al., 2020): a traditional crop which has experienced a low degree of research and commercial interest, due to its smaller seeds and lower yields, compared to major economic crops. One of the only germination-based growth tests for fonio was undertaken by Portères (1955), who observed the developmental stages of germinating fonio seedlings at a range of temperatures up to ∼30°C, observing an optimum temperature of around ∼30°C, and predicting a ceiling temperature of between 40-45°C. Portères commented that these temperatures were perfectly suited to the climate averages in West Africa, during the 1950s. For fonio millets more recently there have been few field or laboratory experiments which examine the specific constraints and germination limits under different climate change scenarios. Senegalese field trials by Gueye et al. (2015) throughout 2010 and 2011 evaluated white fonio (*D. exilis*) sowing and harvesting times versus climate conditions, and Petri-dish germination trials by Iwuala et al. (2021) evaluated the drought response of the close relative and sister crop black fonio (*Digitaria iburua* Stapf.) on three genotypes from the National Cereals Research Institute in Nigeria.

More recently, Pudasaini et al. (2025) conducted an extensive lab-based experiment comparing the difference in morphological response between one white fonio genotype from Mali (arid) and one from Guinea (wet), presenting responses in inflorescence, root and leaf development during irrigated versus drought conditions.

### Modelling future climates and crop suitability

As well as identifying the environmental limits that constrain fonio seed germination, it is useful to compare these with predictions of the temperatures across West Africa in the present and future. Common methodology for predicting what future environments will look like use ‘Shared Socio-economic Pathways’ (SSPs) - a set of 5 possible economic, political, and social development factors (Van Vuuren et al., 2017; Kriegler et al., 2012). These are categorised as: 1, an almost total transition to green energy, and commitment to global development and reducing inequality; 2, little shift from current behaviours and trends; 3, international division which prioritises focus on local regions, with some inequality; 4, increased inequality both within and between regions; and 5, almost total reliance on fossil-fuels and geo-engineering. The SSPs are commonly combined with Representative Concentration Pathways (RCPs) which predict future greenhouse gas emissions, ranging from RCP2.6 with low-emissions, to RCP8.5 - high emissions (Meinshausen et al. 2011). The combination of these two pathway strategies (for example SSP1 and RCP2.6 = SSP126) are then used to predict effects on the environment under different future scenarios: SSP126, SSP245, SSP370, and SSP585.

Every species has its own set of biological limits for living and reproducing, including cereal crops such as fonio. These observed thresholds can be inferred from current geographic occurrences and matched with current climate variables, and future suitable distributions predicted using future climate modelling – which is especially relevant in the scope of exploring how agricultural crops will be affected by climate change. Although this method is entirely theoretical, it can provide a valuable foundation for further interpretation (Pearson and Dawson, 2003), especially when compared to a species’ observed physiological limitations. There are several popular analytical methods which utilise a machine-learning approach to model and predict these species suitability niches in the present and future, including MaxEnt (Phillips and Dudík, 2008; Merow et al., 2013), random forest (Evans et al., 2010), and CLIMEX (Sutherst et al., 1999). Producing an accurate set of results using these methods is dependent on careful selection of data inputs, including valid species distribution data and a set of bioclimatic variables which are relevant both to the species suitability limits, and the geographic region itself (Phillips and Dudík, 2008; Merow et al, 2013).

Fonio was included in a similar study of African crop suitability predictions under future climate change, by Chemura et al. (2024), which uses a combination of climate variables alongside known crop traits, to predict environmental niche suitability under different climate scenarios. Similarly, a plant growth study by Gueye et al. (2013) did not use climate modelling against the raw physiological experiments directly for white fonio varieties in Senegal, but the region-specific scope was useful for predicting the best sowing times in different areas, throughout the year. A combination of these approaches is likely to be effective: seed growth experiments and comparison of thresholds against future temperature predictions, and future environmental niche modelling based on current occurrences.

### Aims and Scope

For this study, as our interest is in determining the thresholds of heat and water potential resilience of fonio for comparison with climates in the future, it is important to test a wide range of accessions from different geographic areas, using a wide range of experimental conditions to detect specific tolerances. We combine two main methods to answer this question: germination trials to determine temperature and water potential thresholds, compared to current and future predicted temperatures in West Africa; and environmental niche analysis using known fonio distribution data (from GBIF) and associated bioclimatic variables, to model future suitability under different SSP scenarios. We also hypothesise that accessions may have different regional tolerances due to its large geographical range.

We investigated the germination thresholds of fonio, for temperature and PEG-induced drought, compare them in the current and future climates, and applied predictive modelling and projections into the near future to assess how suitable fonio cultivation systems are across different areas of West Africa. Here, we report germination profiles of seed accessions from across West Africa, at a range of constant temperatures, and niche suitability models for 2024-2060 under three different SSP scenarios.

## Methods

### Germplasm accessions

Nineteen seed accessions of *Digitaria exilis* for conducting germination trials were accessed from the collections of the Research Institute for Development (IRD) in Montpellier, France, collected up to 50 years ago by partners across Mali, Guinea, Burkina Faso and Togo, and used in the genomic study of Abrouk et al. (2020). Eighteen accessions from across the Fouta Djallon region of Guinea were also accessed from the Herbier National De Guinée (HNG), collected during a study by Burton et al. (2024). Details of all collections can be seen in **Figure** 1 and **Table** 1 – which range from an altitude of 28-1062m, maximum average temperature of 30-43°C, and minimum average temperature of 17-29°C. Accessions 1-19 (Table 1) of white fonio from 4 countries provided an ‘international’ range from two typically separate climates (high maximum average temperature and arid climate in Mali and Burkina Faso, and a lower maximum average temperature and sub-tropical climate in Guinea and Togo; we refer to these as ‘hot’ and ‘cool’ climates). These samples were chosen as a good representation across the wider geographical cultivation region for white fonio, and among genetic populations (the three genetic clusters first identified by Adoukonou-Sagbadja et al., 2007). Accessions 18-37 from the HNG, collected by Burton et al. (2024), were used to provide a wider range of fonio germplasm, to determine average and maximum germination indices.

**Figure 1.**
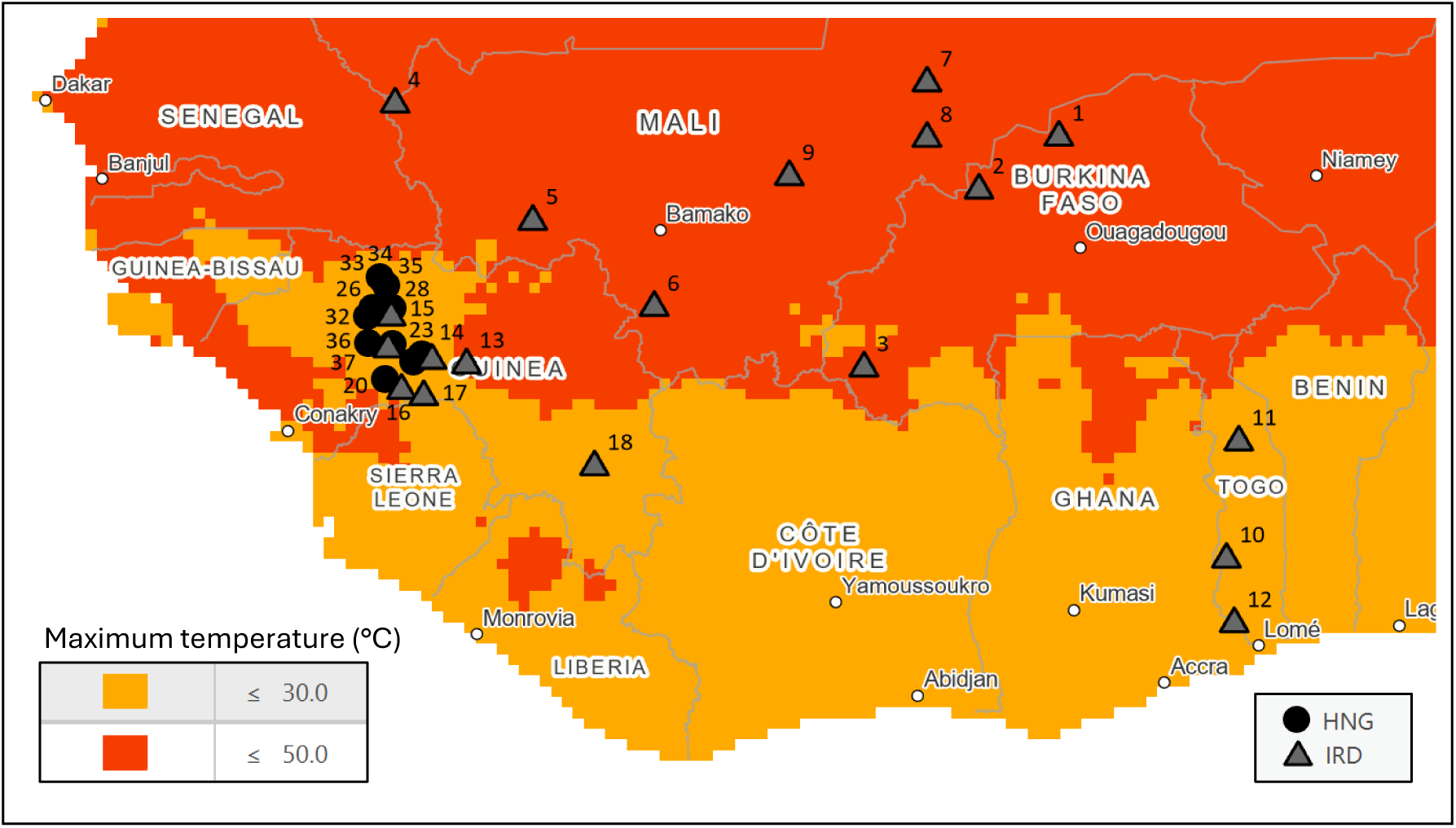
Germplasm collections used in seed germination experiments. Separate collections donated by the IRD (France) and HNG (Guinea) seedbanks are shown, displayed as triangles and circles respectively, and numbered according to accession number given in Table 1. Colours show recent (1970-2000) average maximum temperature across the region in July, separated for above and below 30°C. Accessions collected in Guinea and Togo are classified to be from ‘cool’ vs. ‘hot’ climates in Mali and Burkina Faso.

**Table 1.**
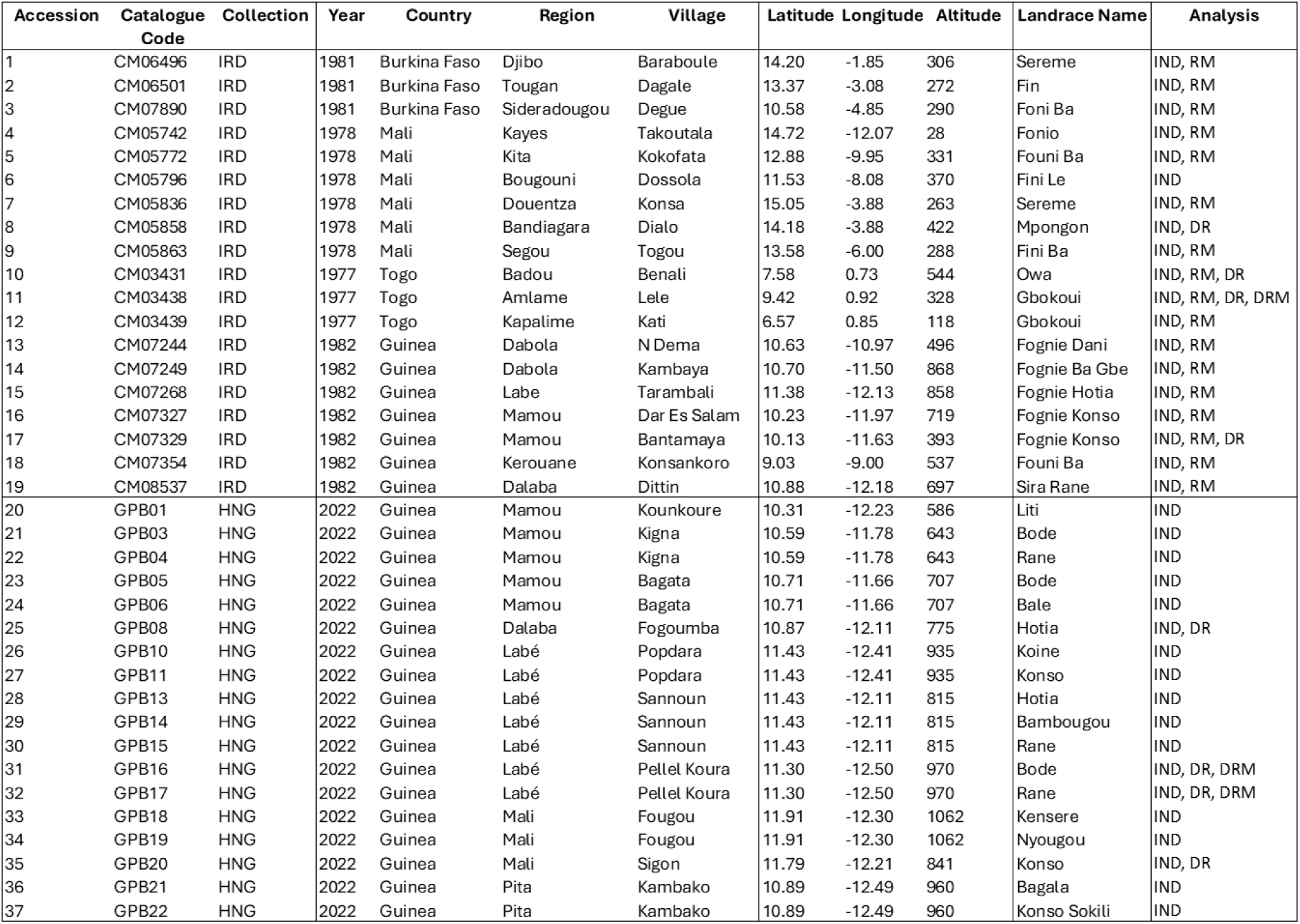
Fonio germplasm accessions used in seed germination experiments as part of this study. Details provided include seed accession numbers used in text (1-37) used in text, catalogue code, and seedbank collection (IRD is the Institut de Recherche pour le Développement, France, and HNG is the Herbier National de Guinée). Analysis column shows which accessions are used for each analysis: germination indices (IND), rate modelling (RM), drought indices (DR) and drought modelling (DRM).

**Table 2.**
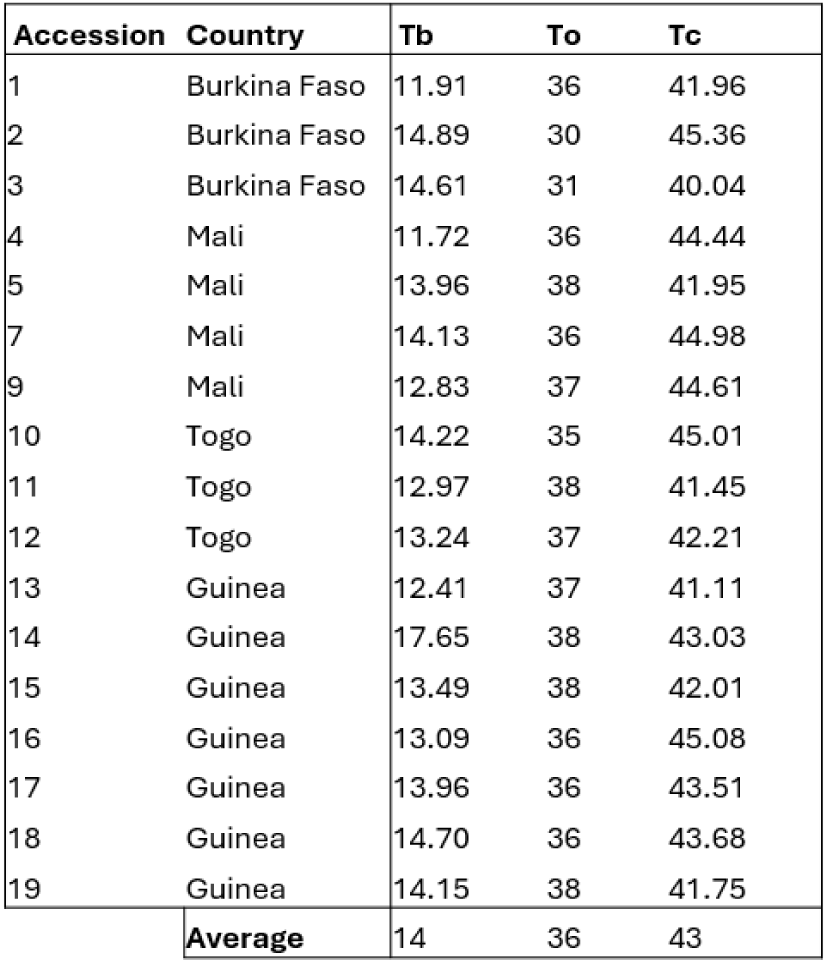
Cardinal temperatures of seed accessions tested as part of germination trials, including base (Tb), optimum (To), and ceiling (Tc) temperatures calculated at t50 (time taken to achieve 50% germination), given in °C. Accessions shown are from the IRD seedbank collection, included as part of an international group also used for statistical analysis against climatic variables from locality of origin.

| Accession | Country | Tb | To | Tc |
| --- | --- | --- | --- | --- |
| 1 | Burkina Faso | 11.91 | 36 | 41.96 |
| 2 | Burkina Faso | 14.89 | 30 | 45.36 |
| 3 | Burkina Faso | 14.61 | 31 | 40.04 |
| 4 | Mali | 11.72 | 36 | 44.44 |
| 5 | Mali | 13.96 | 38 | 41.95 |
| 7 | Mali | 14.13 | 36 | 44.98 |
| 9 | Mali | 12.83 | 37 | 44.61 |
| 10 | Togo | 14.22 | 35 | 45.01 |
| 11 | Togo | 12.97 | 38 | 41.45 |
| 12 | Togo | 13.24 | 37 | 42.21 |
| 13 | Guinea | 12.41 | 37 | 41.11 |
| 14 | Guinea | 17.65 | 38 | 43.03 |
| 15 | Guinea | 13.49 | 38 | 42.01 |
| 16 | Guinea | 13.09 | 36 | 45.08 |
| 17 | Guinea | 13.96 | 36 | 43.51 |
| 18 | Guinea | 14.70 | 36 | 43.68 |
| 19 | Guinea | 14.15 | 38 | 41.75 |
| Average |  | 14 | 36 | 43 |

### Germination trials: temperature and drought simulation

Germination experiments were conducted at the Millenium Seed Bank, Wakehurst, UK, in 2023 and 2024. To test the effect of different temperatures on germination, 20 seeds from each of the 37 accessions were sown in seven 90mm diameter Petri dishes containing water-agar (30ml of 1% Agar, 80-100 Mesh, Fisher Scientific, UK), and placed in incubators (LMS LTD, UK) between 15-45°C at 5°C intervals (constant temperatures at 15°C, 20°C, 25°C, 30°C, 35°C, 40°C, and 45°C), with 3 replicates of each dish. Two photoperiods were applied, 12 hours each of light (LED 55 µmol/m^2^/s) and darkness. 420 seeds and 21 Petri dishes were tested for each accession, at a total of 777 Petri dishes, and 15,540 seeds. Seed germination was scored every 24hrs for 7 days. A germination event was categorised as a radicle over 2mm in length. Trials were conducted in 4 batches, containing a random order of 8-10 accessions in each (between 168-210 Petri dishes per batch). Ungerminated seeds were dissected and classed as viable or empty determined by presence of an embryo, and germination data adjusted accordingly.

To test the effect of drought (changes in water potential) on seed germination, drought conditions were simulated using 6 different concentrations of PEG8000 combined with water, to emulate water potentials at of 0MPa, -0.8MPa, -1MPa, -1.2MPa, -1.4MPa and -1.6MPa. The solutions were prepared using ratios suggested by Michel (1983) and used in similar germination drought experiments by Jadav et al. (2016) and Violita and Azhari (2021): 1ml water containing 0g, 0.25g, 0.3g, 0.32g, 0.34g, and 0.38g, to achieve the respective water potentials. Seeds were germinated on filter paper (Whatman filter paper, 90mm diameter, 200μm thickness) inside a 90mm Petri dish, covered with 3ml of the water:PEG8000 solution. Eight different accessions were tested (8, 10, 11, 17, 25, 31, 32, and 35) which were randomly chosen, but restricted by availability of seed, particularly accessions from Mali and Burkina Faso. Twenty seeds were sown in each Petri dish, across the 8 different water potentials, with 3 replicates for each. Seeds were germinated in incubators at the sub-optimal (20°C) and optimal (30°C) temperatures (determined by temperature trials), and scored every 48 hours for 10 days. Experiments were split into 2 batches, with accessions randomly ordered in each.

### Germination analysis

Germination events for all experiments were recorded per Petri-dish (each given a unique identifying code). This included accession number, repeat, temperature, water potential, viable seed number, number of germination events per time interval (7 time points for heat tests, and 5 for drought tests), and final germination percentage. Final datasets for heat and drought tests were transformed into ‘wide’ and ‘long’ formats (sorted by scoring intervals in the first, and by cumulative germination in the second), following the methods in Onofri et al. (2022). Analysis of germination thresholds, proportion, and rate was conducted in R environment version 4.4.0 (R Core Team, 2024). The package GerminaR (Lozano-Isla et al., 2019) was used to produce germination indices including germination percentage (grp), total number of germinated seeds (grs), mean germination time (mgt), and germination synchronicity (syn)) for each individual petri-dish. These germination indices were analysed using GLM (generalised linear mixed), and one and two-way ANOVA linear models, and conducted separately on each international and regional dataset, following Gianinetti (2020), to examine the effects of temperature, accession, seed age, and collection group. This included data from all 37 accessions. Accessions used in each specific analysis are shown in **Table** 1. T-tests and Bartlett tests were used to analyse the difference between germination of the two climatic groups of accessions. The package DRCTE (Onofri et al., 2022) was used to model germination thresholds, germination rate, and cardinal temperatures for each of the 19 international accessions from the IRD collection, following Onofri et al. (2022) and Catara (2016). A DRCTE (dose-responsive curve time-to-event) model object was produced using the NPLME method, and germination rate as a response to temperature across time was calculated using an exponential model (GRT.Ex) in the drcSeedGerm package (Onofri et al., 2018). Thermal thresholds were modelled at time taken for a Petri-dish to reach 50% germination (t50). The predict() function in R was used to calculate optimum temperature (To) for each accession using data from the GRT.Ex model. Seven accessions where models would not converge above t40 were excluded from this analysis: 6, 8, 20-24, and 37.

Similar methods were used to analyse data from drought experiments. Data from trials at 20°C and 30°C were separated, and germination indices calculated for all 8 accessions tested (details in **Figure** 1). Germination rate curves were calculated using the water potential model GRPsiPol (rather than GRT.Ex) in drcSeedGerm to explore the relationship between water potential and germination rate. t50 was used for model convergence at 30°C, and t15 at 20°C; only 3 accessions (11, 31 and 32) could be modelled to t50 at 30°C, and none at 20°C. At t15 and 20°C only 3 accessions (10, 31, and 35) could be modelled.

### Phenotypic data

After seed germination experiments were complete (at the end of 7 days for heat and 10 days for drought), the weight of total leaf matter from 5 individual seedlings from 3 Petri-dishes tested at 20°C, 30°C, and 40°C was measured in milligrams (mg) using a microscale (leaves from a total of 45 seedlings per accession across treatments), for 17 accessions (1, 3, 8, 10, 11, 13, 16, 17, 21, 24, 25, 27, 28, 31, 32, 35, and 37). Seedlings grown as part of drought trials were also weighed at 0MPa and -1MPa, accessions 10, 25, 32, and 35 grown at 30°C.

Seedlings from accession 1 were photographed periodically over 7 days to capture the development of the seedling during heat and drought experiments to record qualitive differences across the gradient of conditions.

### Climate Data and Niche Modelling

Climate data was accessed and downloaded from WorldClim ver. 2.1 (https://www.worldclim.org/), all at 10min resolution. For present/historic climate data we used average maximum temperature (°C) (tmax) from the years 1970-2000, for the months 5, 6, 7, and 8 (May, June, July, August) to represent the temperatures experienced by seeds germinating during the sowing period (Burton et al., 2024). This was compared to the maximum temperature of the same months in future climate predictions, for the 2040-2060 period, under the SSP126, SSP245, SSP370, and SSP585 scenarios, using the CMCC-ESM2 (Euro-Mediterranean Center on Climate Change Earth System Model) general circulation model (GCM). Climate values were attached to xy coordinates and averaged across the sowing period, to predict a count of occurrences occurring within different temperature ranges shown to be suitable for successful germination (<35°C).

For modelling crop suitability under future climates, occurrence data for *Digitaria exilis* was downloaded from GBIF (https://www.gbif.org/species/5289953) as representative of the present occurrence range of the species. GBIF records did not include any occurrence data from Senegal or Nigeria, which are known to be key cultivation regions for white fonio (Cruz et al. 2016), and verified records with co-ordinates from these regions could not be accessed. These absent occurrences were instead used to assess the validity of the suitability models (i.e. see whether the model still predicted these areas to be suitable, despite missing data). Bioclimatic variables for the present (1970-2000) and future (2040-2060) were downloaded from WorldClim (https://www.worldclim.org/data/index.html), under the SSP126, SSP245, SSP370, and SSP585 scenarios, using the CMCC-ESM2 general circulation model (GCM). A full range of all bioclimatic variables were input to niche modelling analysis first, and the variables with the lowest model contributions removed systematically. The remaining bioclimatic variables BIO3 (Isothermality), BIO4 (Temperature Seasonality), BIO5 (Max Temperature of Warmest Month), BIO9 (Mean Temperature of Driest Quarter), BIO10 (Mean Temperature of Warmest Quarter), BIO14 (Precipitation of Wettest Month), BIO15 (Precipitation Seasonality), and BIO17 (Precipitation of Driest Quarter) were chosen as the most useful climatic variables which are likely to be significant indicators of fonio suitability.

Raster layers and maps were processed using ArcGIS Pro 3.4.0. The maps containing current and future variables were cropped to the extent of West Africa, and raster layers converted into ascii files. The current variables and species distribution data were input to Maxent 3.4.3 (Phillips and Dudík, 2004, available from https://biodiversityinformatics.amnh.org/open_source/maxent/), and run to produce an initial map of current species suitability, using the Cloglog model. This was rerun with random test percentage at 30%, which removed a random 30% of the occurrences, to test likelihood with a smaller dataset for the relaxed minimum training presence and strict 10^th^ percentile training presence. The full model was then rerun again with the future climatic variables as projection layers, which was repeated for each future scenario. These results were overlaid in ArcGIS Pro, and maps of regions suitable in the present versus future produced using the Raster Calculator tool and binarized suitable niche maps. Percentage of suitable area was calculated using raw count data provided in ArcGIS.

## Results

### Response to temperature

Germination synchronicity between accessions (syn), which measures the consistent overlap of germination within groups, is found to be insignificantly affected by temperature and accession (p>0.05). Neither date of seed collection nor collection group (IRD vs. HNG) have a significant effect on seed germination time, germination proportion, or synchronicity (p>0.05; **Figure** S1).

Temperature had a significant effect on seed germination percentage across all accessions (p<0.005) when analysed using a binomial GLM including accession and temperature as fixed effect dependent variables, and grp as a response. A set of graphs demonstrating the relationship between heat and water potential, and seed germination indices and leaf weight, are given in **Figure** 2. The temperature with the mean highest successful germination was 30°C (87.2%), followed by 25°C (86.8%), and 35°C (86%). No germination was observed at 45°C, confirming that this temperature exceeds fonio’s possible germination range. A significant relationship between mean germination time (mgt) and temperature was also observed (p<0.005), where the temperature with the fastest germination was 35°C (1.59), followed by 40°C (1.79), and 30°C (1.85), measured in average time taken for germination.

**Figure 2.**
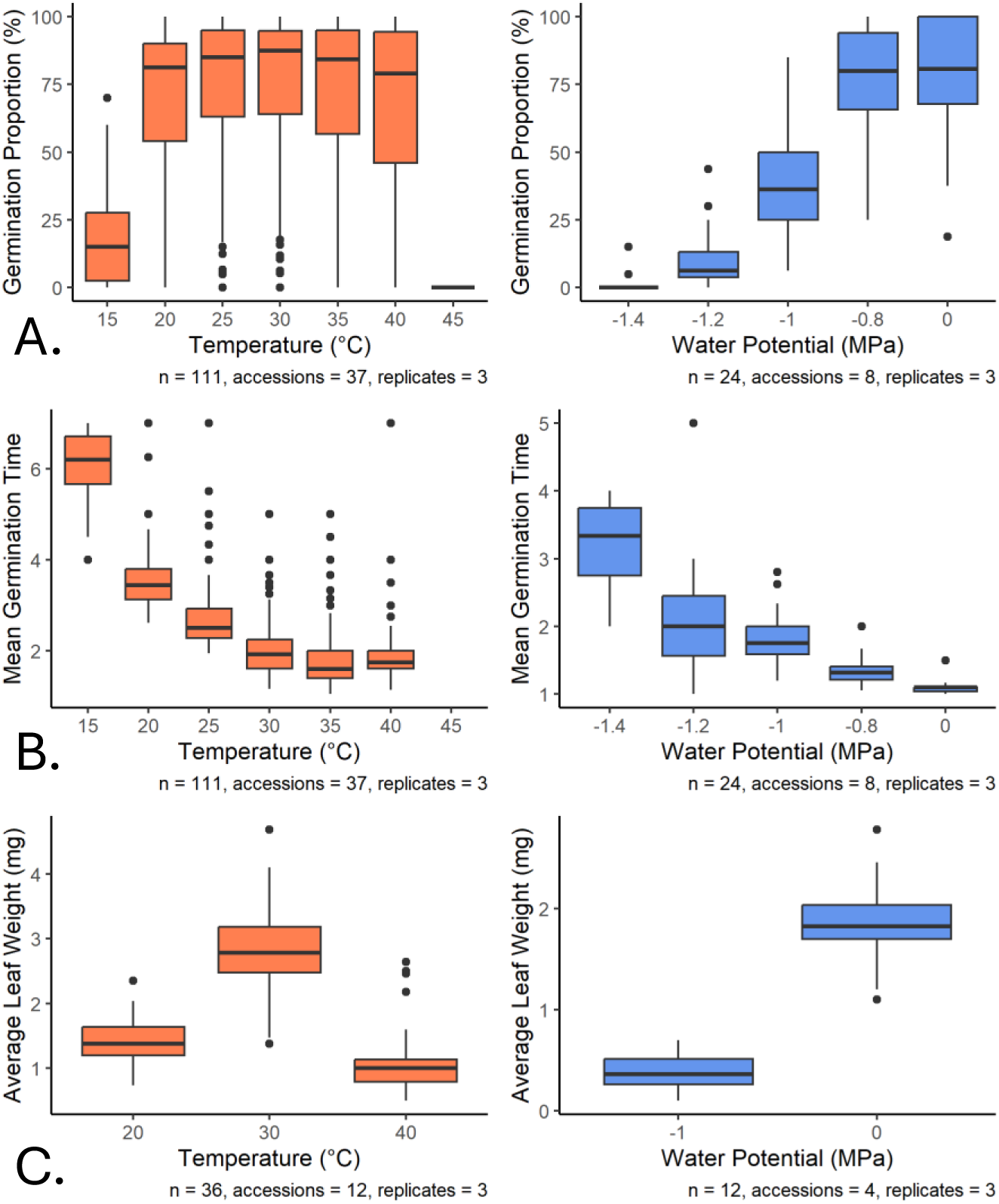
Response of seed germination proportion (row A), mean germination time (row B), and leaf weight (row C), to temperature (boxplots in red) and water potential (boxplots in blue). Graph captions give sample size (n) for each boxplot, and number of seed accessions used. MGT measurement is given as average time interval taken for successful germination.

Cardinal temperatures were modelled for 19 accessions using t50 (time taken to achieve 50% successful germination per Petri-dish). Average Tb was found to be 14.52°C (SD=1.7°C, maximum 17.99°C, minimum 11.72°C), To was 36°C (SD=2.15°C, maximum 38°C, minimum 30°C), Tc was 42.62°C (SD=1.68°C, maximum 45.36°C, minimum 39.99°C) and mean thermal window (Tc-Tb) of possible germination found to be 28°C (SD=2.65°C, maximum 32.72°C, minimum 23.73°C), and all were significant at p<0.001.

Estimated values for cardinal temperatures are shown in **Table** 2, and curve for germination rate and cardinal temperatures for accession GB31 in **Figure** 3. A full set of results for both temperature and water potential experiments are given in **Figure** S2, and germination rates and proportions for all seed accessions under heat conditions are shown in **Figure** S3.

**Figure 3.**
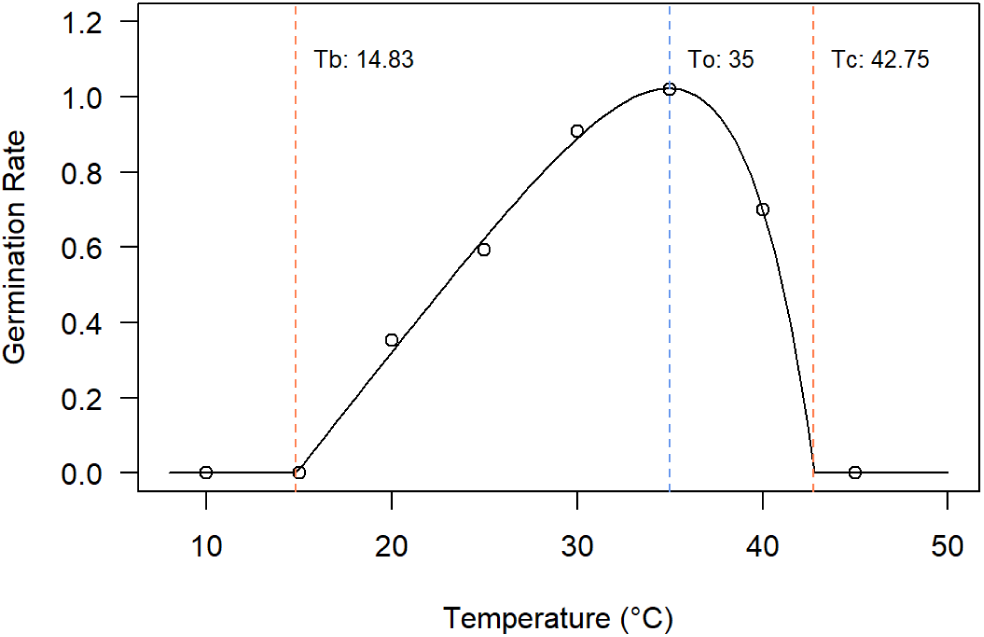
Modelled germination rate for accession 31 from Guinea, under different temperature conditions, at t50. Rate is given as a log-adjusted value based on average time taken for all seeds in a Petri-dish to achieve 50% germination. Dashed red lines show viable germination limits at base (Tb) and ceiling (Tc) temperatures, and blue line shows predicted optimum temperature (To).

Correlations between average climate data associated with accession collection localities (maximum temperature, minimum temperature, and average precipitation) and the climate groupings assigned to each accession (hot and arid vs. wet and sub-tropical) are significant, and the relationship between climate (arid Burkina Faso and Mali grouped together versus Guinea and Togo) and seed germination percentage, at the temperatures of 30°C and 40°C were all found to be significant (p=0.008 and p=0.0005, respectively). However, there is significantly unbalanced variance within groups (Bartlett test p<0.005, F=0.298, F=0.254, and F=0.149. This is shown in **Figure** S4, with a full table of statistical correlation between all variables in **Figure** S5. There was no correlation between Tb or Tc with country or climate groups. There is a similarly minor significant relationship between To and both country and climate group, (p=0.0009284 and p=2.088e-09, respectively), though again there is significantly unbalanced variance between groups (Bartlett test p<0.005, F=0.132).

### Response to drought

Germination synchronicity was significantly affected by water potential (p<0.001). Water potential (WP) had a significant effect on seed germination percentage (p<0.001), using a one-way ANOVA linear model. The mean highest successful germination percentage at 30°C was at 0MPa (77.1%), followed by -0.8MPa (74.8%), -1MPa (38.4%), -1.2MPa (10.5%), and -1.4MPa (1.25%). Mean germination time was also significantly affected by WP (p<0.001), where again the fastest mean time to germination was found at 0MPa (1.09), followed by -0.8MPa (1.33), -1MPa (1.80), -1.2MPa (2.12), and -1.4MPa (3.16). At 20°C the mean seed germination was highest again at 0MPa (75.6%), followed by -0.8MPa (5.05%). There was no germination at -1MPa to -1.4MPa. Germination time was also lowest at 0MPa (1.71) versus at -0.8MPa (3.64).

The minimum water potential required for germination (Ψb), by rate, could only be modelled for 3 accessions to t50 at 30°C. Average Ψb was -1.09MPa, with a minimum of -1.22MPa (accession 11 from Togo), and a maximum of -1.01MPa (accession 32 from Guinea). Germination rate curves for water potential experiments are shown in **Figure** S6. Germination at 20°C could only be modelled using t15, at which the mean Ψb is -0.8MPa, from three accessions.

### Seedling phenotypes of stress

Temperature had a significant effect on seedling weight (p<0.001), the highest mean weight was found at 30°C (2.81mg), followed by 20°C (1.41mg), and 40°C (1.11mg). This was independent of accession (p<0.001). Water potential also had a significant effect on seedling weight (p<0.001), with the highest mean weight of 1.87mg at 0MPa and 0.4mg at -1MPa, also unaffected by accession (p<0.001). Mean weights of leaves are shown in **Figure** 2.

Observable phenotypes of temperature conditions at 7 days after sowing include: undeveloped roots and shoots with no clear coleoptile expansion at 15°C; coiled, non-branching roots with a single leaf at 20-25°C; roots with multiple branchings and multiple leaves at 30-35°C; and long unbranched roots, elongated and pallid leaves at 40°C. Phenotypes of temperature and drought stress shown in **Figure** 4.

**Figure 4.**
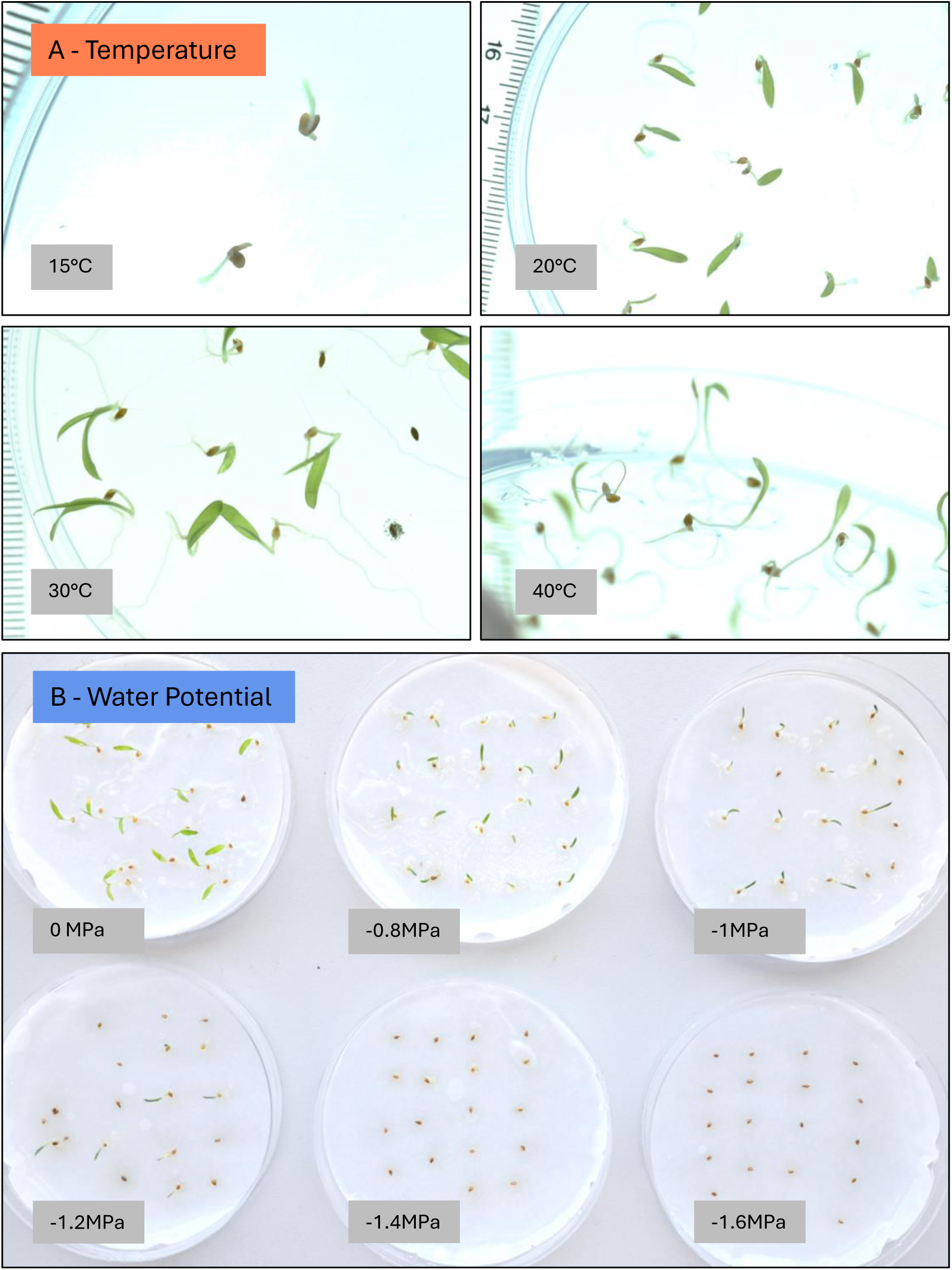
Seedling growth phenotypes of seed accession 1 from Burkina Faso, 7 days after sowing, germinated under different heat (A) and water potential (B) conditions. Germinated seed can be seen at -1.4MPa in B, second seed from the right in second row from the top.

### Germination temperature thresholds vs. future temperatures

Historic (1970-2000) and future (2024-2060) projected maximum temperature values across West Africa for the main fonio sowing period (May-August) are shown in **Figure** 5. These include future projections under 4 different scenarios, and temperature categorised to match seed germination testing thresholds. The threshold of above 35°C, corresponding to average To, is chosen as unsuitable for consistent successful germination, based on seed germination results and seedling phenotypes described above. Large parts of Senegal, Burkina Faso, and Mali experience average maximum temperature of between 35-45°C in May and June in current climates – in future climates this range increases southward. In July, only fonio cultivated to the northern most extent of range (in Mali) experience temperatures of 35-45°C in the present, but this range extends to cover large parts of cultivation in Mali and Burkina Faso in the future, especially under SSP585. August remains within the window of likely germinable temperatures in current and all future scenarios. When occurrence points are matched to maximum temperatures under each current and future scenario, out of 1609 data points, and averaged across all 4 months in current conditions, 88.3% of occurrences are within a suitable temperature range. This decreases to 79.86% to 79.08% under SSP126, SSP245, and SSP370, and down to 71.05% for SSP585. This represents a decrease of 10.45% between current and future (SSP126-370) scenarios - shown in **Figure** 6, and separated by month in **Figure** S7.

**Figure 5.**
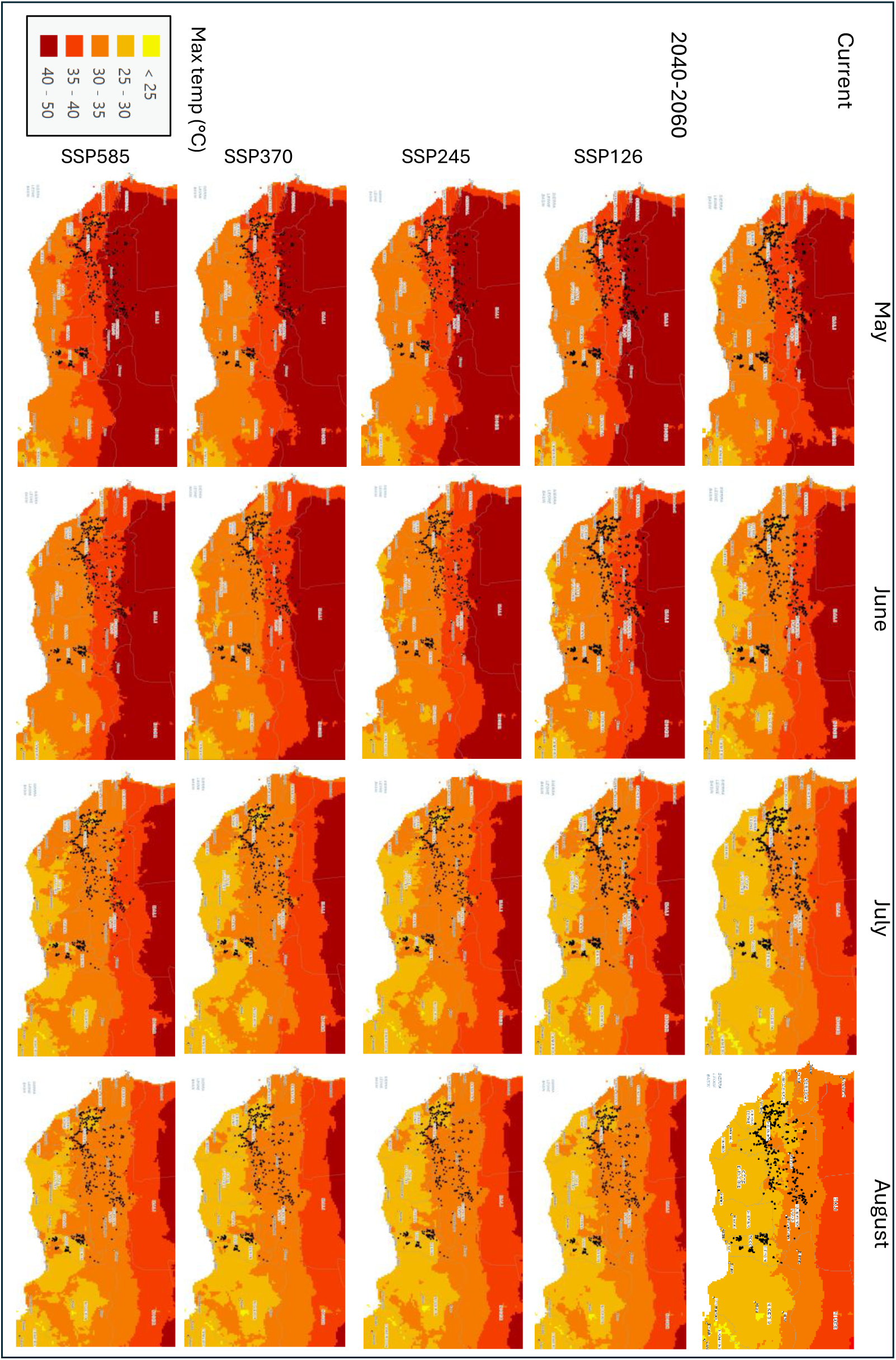
Current and future average maximum temperatures across West Africa, in the fonio sowing months from May to August. Future predictions are shown for 4 future SSP scenarios. Black circles show fonio occurrences recorded on GBIF (https://www.gbif.org/species/5289953). Areas are coloured by maximum temperature range from yellow to dark red – areas in red and dark red (Temperatures >35°C are above the average optimum fonio germination temperature.

**Figure 6.**
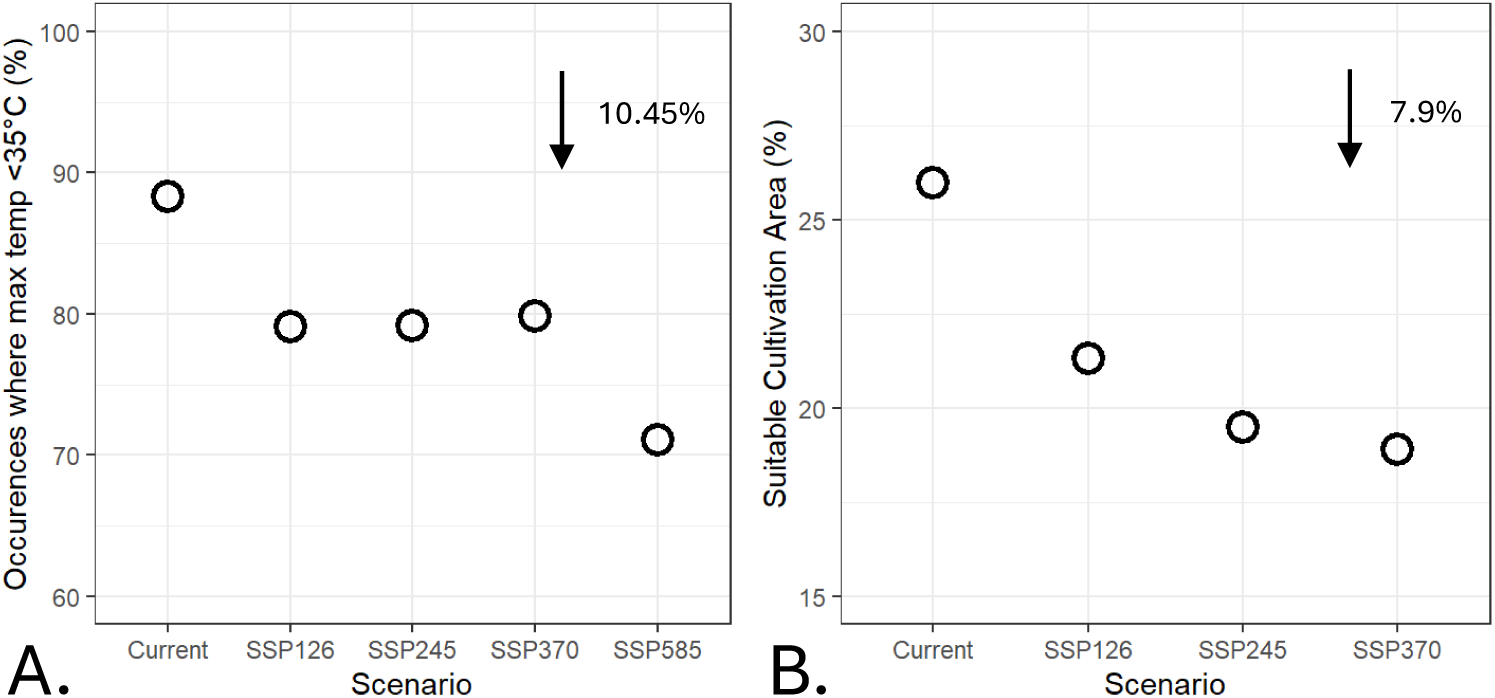
Suitability of fonio cultivation in current and future climate scenarios, calculated by two methods: A. occurrences in geographic areas below the optimal temperature threshold of 35°C; and B. niche suitability modelling. Arrows and labels show the decrease in mean suitable occurrences and area between current and future scenarios.

### Niche suitability modelling

Niche suitability climate maps were produced under current and 3 future (SSP126, SSP245, SSP370) scenarios. Based on 30% test data and a minimum training threshold, the omission rate was 0 (binomial test p<0.050. AUC based on 30% removal of test data = 0.905. The minimum training threshold is 0.041, and the 10^th^ percentile training presence is 0.272, equal training sensitivity and specificity is 0.389. Fractional predicted area at the 10^th^ percentile training presence is 0.239 (24%). All of these are significant to p<0.0001. The highest percentage contribution of bioclimatic values to the model was temperature isothermality (BIO4, 58%), followed by precipitation seasonality (BIO15, 13.3%), precipitation of driest month (BIO14, 9.4%) and mean temperature of driest quarter (BIO9, 7.2%). The lowest contributor was mean temperature of warmest quarter (BIO10, 1.2%).

Suitability maps for fonio are shown in **Figure** 7. In current and recent historic conditions, 25.9% of the whole region shown is predicted to be suitable for fonio cultivation. This area covers nearly all fonio occurrences on the map, including Senegal and Nigeria, which are known to cultivate fonio, but no available occurrence points could be used for this study. This suggests an accurate modelling of crop suitability in the current climate. Under predictions in SSP126 the percentage of suitable area decreases to 21.3%, to 19.4% at SSP245, and to 18.8% at SSP370. This represents a range of between 6.3% to 9.1% reduction between current to future, and an average of 7.9%, shown in **Figure** 6. Based on the 10min resolution raster files used, this represents around 400,000km^2^ (average of 1190 squares in raster layer missing in future) of land. There is also an increase of some areas newly suitable for fonio, especially in Sierra Leone, Cote D’Ivoire, Ghana, and Nigeria, as the southern regions of West Africa experience higher average temperatures – a total average increase in new areas of 5.5% (around 250,000km^2^).

**Figure 7.**
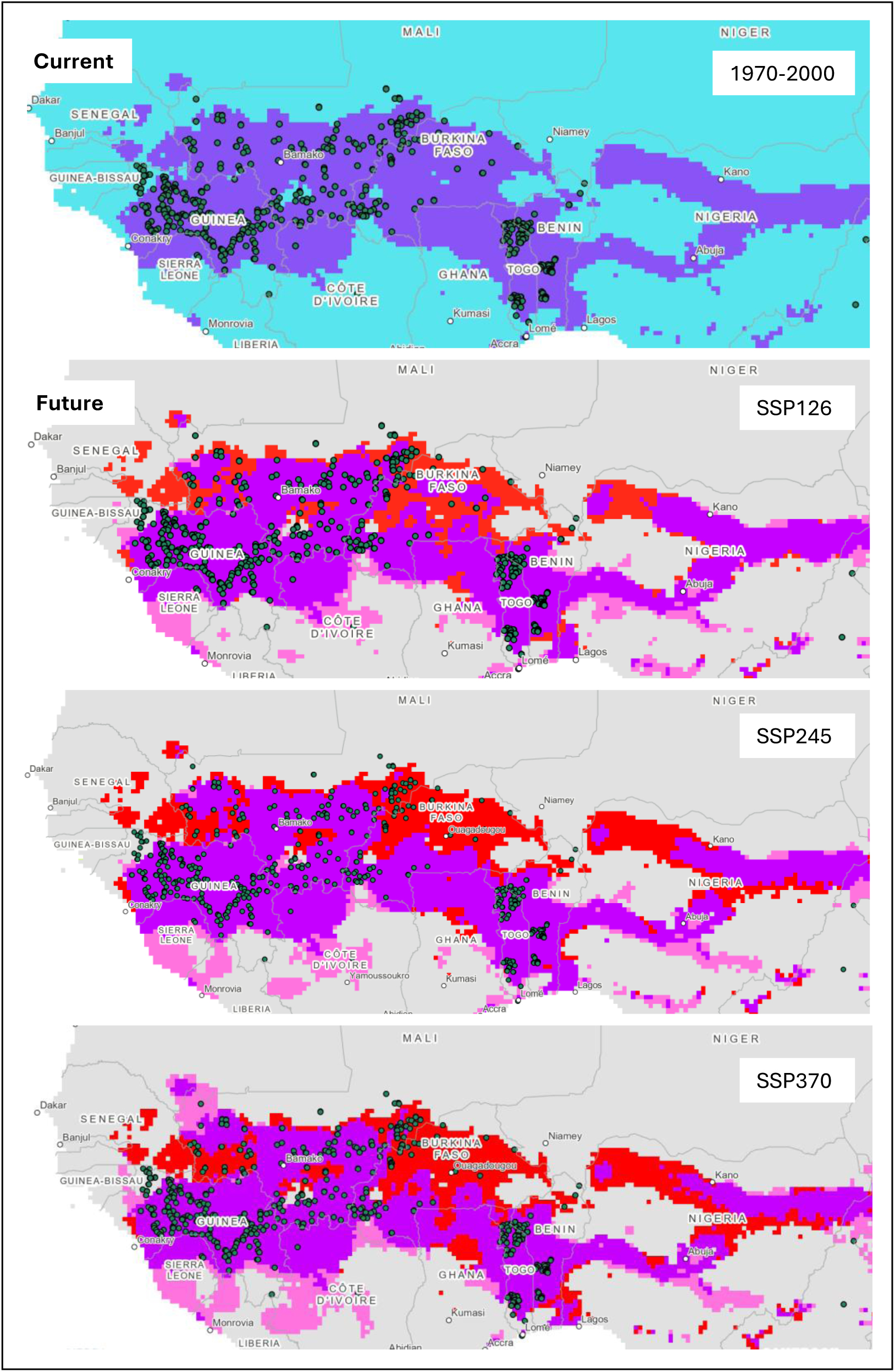
Suitable area for fonio cultivation under current and future projections, calculated using environmental niche modelling. Fonio occurrences from GBIF are shown as circles in green. Area in blue shows unsuitable area in present, purple shows suitable area both in the current and future, red shows suitable area missing in future projections, and pink shows newly suitable area.

## Discussion

### Germination responses to climate change

The germinable temperature thresholds of *Digitaria exilis* were found to be between an average Tb of 14.52°C (observed down to 11.72°C) up to an average To of 36°C. Germination rate models estimated the highest Tc to be 45.36°C, though no seed tested in germination trials was able to germinate successfully at 45°C. This corroborates the results of Portères (1955), whose germination experiments suggested that fonio has a Tc of below 45°C, and a To of above 30°C.

The effects of seed age (collected in1977, 1978, 1981 and 1982 vs. 2022), and country and climate of origin (Mali and Burkina Faso vs. Guinea and Togo), appeared to be negligible. In traditional storage, the dried grains are reported to be germinable for several years after harvest (Cruz et al. 2016), but it appears that there is little to no difference in germination success between seeds harvested, dried and stored in the 1970s, and those stored and dried 50 years later, and no observable dormancy. Many orthodox plant seeds remain germinable after long periods, although it is uncommon to not see a significant decrease in germination success after such a long period of time (Walters et al., 2005; Kameswara Rao et al., 2017; De Vitis et al., 2020). This relationship between seed age and germinability is obviously very dependent on storage methods, and highlights the positive role of seed-banking in preserving valuable indigenous crop seeds for growth in the future.

Although country and climate groupings had a minor statistically significant relationship with seed germination proportion (grp) and optimal temperature (To), there was disproportionate variation between groups, where seeds from arid environments (Mali and Burkina Faso) had a more varied germination response to higher temperatures, and actually fail to outperform seeds from cooler climates found in Guinea and Togo. There appears to be no definite difference between germination responses between the two groups that would suggest strong adaptations to local climates. Generally, there is little variation in germination response to high heat among varieties of fonio, even those from vastly different geographic areas. This may be due to our experimental conditions and climate variables not taking into account the specific microclimates that older generations of seed experienced, or that sample size was not large enough to capture any strong, consistent difference between groups.

It is also known that fonio varieties are often transported to different regions through trade to neighbouring villages and to different habitats (upland slopes, lowland grasslands; Burton et al., 2024; Cruz et al., 2016). This would suggest a crop that is frequently moved between different nearby microclimates, possibly explaining why a significant difference in origin climate-specific heat tolerance was detected in this study. However, it is more likely that fonio is hitting an absolute limit with germination temperature, where adaptation is no longer feasible due to physiological constraints. In a recent study by Pudasaini et al. (2025), two fonio genotypes, one each from Mali and Guinea, were compared for differences in drought response in whole mature plants. While they found that there was strong difference in root architecture and yield (the variety from Mali outperforming the variety from Guinea), they make no comment on any differences in the overall germination success or early seedling development of the two varieties. This again supports that germination vigour under heat stress is generally high across white fonio varieties, but also reaching a set physiological limit of around 42°C. This high resilience is possibly a trait conserved from the likely origins of *Digitaria* alongside the subtribe Anthephorinae and other Panicoid C4 grasses in arid central and East African grasslands (Gallaher et al., 2022; Bouchenak-Khelladi et al., 2010).

### Suitability of fonio in future climates

From the germination thresholds and phenotypes described above, it is sensible to predict that fonio being sown in any areas that experience consistent average maximum temperatures above 35°C for prolonged periods is likely to experience a decrease in plant health (at 40°C) and germination success and survival (above 40°C), resulting in decreased grain yield and food availability.

Analysis of seed germination results against future climate predictions above utilises a comparison of realistic thresholds for successful fonio seed germination (below 35°C) compared to current and future maximum temperatures across West Africa. In the current climate, very few occurrences appear within the regions which experience >35°C temperatures in July and August, apart from 3 occurrences in the area on the border between the Mopti and Tombouctou regions in Mali, and Burkina Faso. In May and June large areas of the upper Sahel are unsuitable, but most areas of Guinea are suitable throughout the whole season. This supports why farmers in Guinea described how early varieties of fonio may be sown as early as May in a study by Burton et al. (2024), and there has recently been a shift to sowing later seasons, and abandonment of early varieties. Meanwhile, Mali, Senegal, and Burkina Faso appear suitable for these earlier varieties. A growth study by Gueye et al. (2015) with fonio in south-eastern Senegal, suggests that early July had the best sowing dates for plant growth and grain yield, as it avoided the intense heat of May and June, and avoided rains in August. This fully corroborates with our results shown here, and suggests that in the future, farmers across West Africa, especially in arid areas, are likely to have to shift their sowing and harvest calendars to better suit the heat tolerance of their crop seeds to more suitable later months.

Between the current and future climate scenarios, using results from seed germination trials against future climates, around 10% of suitable land for fonio cultivation is likely be lost. The months May and June experience far more severe reductions than July and August, although in some scenarios July is also affected. Key cultivation regions in Senegal, Mali, and Burkina Faso appear to be affected the most, where some areas experience temperatures above 35°C more frequently. These results support those from the niche suitability model predictions: reduction in suitable niche area for fonio cultivation under these models predict a reduction of around 8% across the region. The area in Mali that appears to be currently unsuitable with the 3 occurrences in the first method, are also unsuitable in the current climate under these predictions. Even under the most optimistic scenario SSP126, there is a significant reduction in unsuitable niche area of around 6%, where the model entirely removes Senegal, and large parts of Mali and Burkina Faso from the suitable land area. This only degrades further as the SSP scenarios become decreasingly optimistic. Crop suitability models produced by Chemura et al. (2024) similarly found that fonio by 2050 will experience a 2-7% reduction in suitable area under the different SSP scenarios – though this was calculated to the scale of the whole African continent, their predictions also suggest that the upper Sahel countries will experience the most severe reductions. A review study on the effect of climate change on crop yields in West Africa found a median yield loss of ∼11% in the future, which was far more impactful in northern West Africa (including Mali and Burkina Faso) vs. southern regions of Guinea and Ghana (Roudier et al., 2011). The results in this study for fonio then of around 10% decrease in the future is similar to other studies, tested through multiple different methods. This reduction of suitable area for fonio, coupled with low specific local adaptation for germinating in extreme arid climates, is worrying, considering how viable germination already appears to have reached a consistent limit of around 42°C in seed accessions across the cultivation range.

Three key areas that appear to be affected in both methods are south-eastern Senegal, the south-eastern Mali:Burkina Faso border, and north-eastern Benin, areas shown in **Figure** S8. These areas are historically very important for the cultivation of fonio, with records of fonio in this area of Benin as far back as 900-1400AD, and possibly having been domesticated near the Mopti region in Mali before 500BC (Champion and Fuller, 2018). In a study by Porcuna-Ferrer et al. (2024) on the south-eastern fonio cultivation by Bassari farmers in Senegal, they reported that a mix of socio-political issues and policies promoting non-indigenous crop species over fonio and Bambara groundnut, causing a decline in fonio production of up to 40%. Around 9% of interviewees cited environmental factors as the reason for abandonment, suggesting that cultivation in currently climate-suitable regions across West Africa will and do now experience abandonment – the effects of climate change are also coupled with and intensified by social attitudes of governments and local communities.

The climate change between present and future scenarios are also beneficial in some areas: in Sierra Leone, Cote D’Ivoire, Ghana, and Nigeria, suitable land for fonio is predicted to expand by around 5.5%. A study by Wimalasiri et al. (2023) investigated a similar response for proso millet (*Panicum miliaceum* L.), using the 2024-2069 predictions from the C3MP/CMIP5, with increased yield under future rainfall increases, alongside a 2°C temperature increase in the dry zone of Sri Lanka. Chemura et al. (2024) while predicting an overall decrease in suitable land for fonio, predicted a positive increase in suitable area for similar millet species sorghum and pearl millet, and also Bambara groundnut, by up to 5% by 2050.

## Conclusions

This study provides critical insights into the climatic resilience of fonio. We have established that the physiological growth limit of fonio is up to a likely absolute maximum of 43°C, with an average optimum temperature of around 35°C based on extensive germination experiments involving 37 seed accessions collected from across multiple habitats and climates in West Africa. There appears to be no straightforward relationship between heat resilience in germination trials and the seed accessions’ local climate, suggesting a low selection for heat-resilient varieties, or local adaptation to the climate, and instead a more general high tolerance for heat for the species.

These results highlight both opportunities and risk for fonio as an important indigenous crop. When comparing temperature suitability thresholds to current and future climates using two different methods, we find that over the next 40 years, fonio cultivation in regions close to the Sahel in Senegal, Benin, Mali, and Burkina Faso, are very likely to be adversely impacted by climate change. These are some of the most important current and historical areas for fonio cultivation, both socially and economically. However, many other areas will remain or transition into the optimal climate window for cultivation, including Sierra Leone, Cote D’Ivoire, and Ghana. While fonio remains an essential component of food security in West Africa, the increasing threat of climate change necessitates improved adaptive agronomic strategies, such as shifting sowing times to avoid extreme heat and weather events, accelerated selection and breeding for more heat-tolerant varieties, and expanding cultivation into newly suitable areas.

From this study, future research and policy may be implemented to target conservation for the most under-threat areas, and promote fonio in regions where it is still vitally suitable and sustainable, where other crops are not. Policies that promote the conservation and commercialisation of neglected crops like fonio will be crucial in supporting food security in climate-vulnerable regions of the Sahel. These include policy initiatives like the Comprehensive Africa Agriculture Development Programme (CAADP, 2025) administered by the African Union, which aims to end hunger (Malabo Commitment 3), and enhance resilience of livelihoods and production systems to climate variability (MC6). Our research provides a further opportunity to conduct similar studies on other dryland NUS crops (neglected and underutilised, Ulian et al. 2020) in order to inform agricultural development policies, programmes, and projects on the basis of comparative advantages and disadvantages of climate-resilient crops.

## Supplementary Data

Figure S1. Effect of date collected and seedbank collection group on germination proportion.

Figure S2. Cardinal temperatures (optimum temperature, ceiling temperature, base temperature, and thermal window at t50) and base water potentials (Psib) at 30°C (t50) and 20°C (t15) for fonio seed accessions.

Figure S3. Modelled germination rate curves and proportions (boxplots) for all accessions used in temperature experiments.

Figure S4. Effect of seed accession origin climate (Guinea and Togo vs. Mali and Burkina Faso) on germination proportion at 30°C, 35°C, and 40°C.

Figure S5. Correlation of variables between seed germination proportions, cardinal temperatures, and climatic variables. Significant p values are in bold.

Figure S6. Modelled germination rate of three fonio accessions under a gradient of water potential conditions, at threshold t50.

Figure S7. Percentage of fonio occurrence points mapped to maximum temperatures above 35°C, in the current and under future climate SSP scenarios, grouped by month.

Figure S8. Key areas predicted to experience significant reduction in suitable land for fonio cultivation.

## Acknowledgements

We are extremely grateful to the staff and students at the Herbier National de Guinée including director Sékou Magassouba, and Kew Science liaisons Charlotte Couch and Martin Cheek for helping with fieldwork to collect fonio seed material. We thank Adeline Barnaud at the IRD and Claire Billot at CIRAD (both Montpellier, France), for their valuable collaboration and for providing seed material from the IRD for our study. We also thank the curation, research and laboratory teams at the Millenium Seed Bank (UK), including Pablo Gomez Barreiro, Silvia Bacci, and Bendetta Gori for assisting with germination experiments. Thank you to Owen Corbett for guidance on producing suitability models in ArcGIS. This study is a part of G. Burton’s PhD project, and we are thankful for funding and administrative help provided by Christiane Morgan at the Grantham Institute (Imperial College London), and Felix Forest at Kew Science.

## Author Contributions

GPB-TU: conceptualisation; GPB, HMB, CES, MSV, EM, and TU: methodology; GPB, CES, EM: data analysis; GPB-TU: writing; PC, RMG, PR, MSV, EM, and TU: supervision; GPB, MSV, and TU: funding acquisition.

## Data Availability

Data from this study will be deposited to Dryad online repository after review.

