## Supplementary Material for "Impacts of climate change on fonio millet: seed germination and suitability modelling of an important indigenous West African crop" (from another version of this preprint: version 1, 2025-03-17)

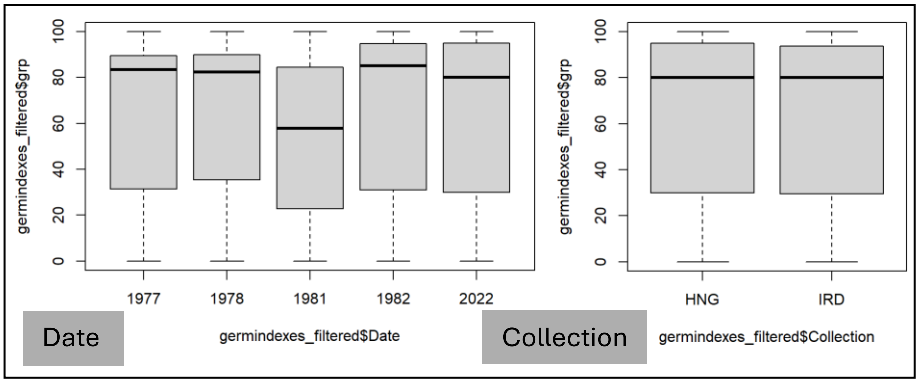


Figure S1 – Effect of date collected and seedbank collection group on germination proportion.

| Code | Collection | Country | Landrace | To | Tb | Tc | Tw | Psib°C 30 | Psib°C 20 |
|---|---|---|---|---|---|---|---|---|---|
| 1 | IRD | Burkina Faso | Sereme | 36 | 11.91 | 41.96 | 30.05 | - | - |
| 2 | IRD | Burkina Faso | Fin | 30 | 14.89 | 45.36 | 30.47 | - | - |
| 3 | IRD | Burkina Faso | Founi Ba | 31 | 14.61 | 40.04 | 25.43 | - | - |
| 4 | IRD | Mali | Fonio | 36 | 11.72 | 44.44 | 32.72 | - | - |
| 5 | IRD | Mali | Founi Ba | 38 | 13.96 | 41.95 | 27.99 | - | - |
| 7 | IRD | Mali | Sereme | 36 | 14.13 | 44.98 | 30.85 | - | - |
| 9 | IRD | Mali | Founi Ba | 37 | 12.83 | 44.61 | 31.78 | - | - |
| 10 | IRD | Togo | Owa | 35 | 14.22 | 45.01 | 30.79 | - | -0.84 |
| 11 | IRD | Togo | Gbokoui | 38 | 12.97 | 41.45 | 28.48 | -1.217 | - |
| 12 | IRD | Togo | Gbokoui | 37 | 13.24 | 42.21 | 28.97 | - | - |
| 13 | IRD | Guinea | Dani (Rane) | 37 | 12.41 | 41.11 | 28.70 | - | - |
| 14 | IRD | Guinea | Ba Gbe | 38 | 17.65 | 43.03 | 25.38 | - | - |
| 15 | IRD | Guinea | Hotia | 38 | 13.49 | 42.01 | 28.52 | - | - |
| 16 | IRD | Guinea | Konso | 36 | 13.09 | 45.08 | 31.99 | - | - |
| 17 | IRD | Guinea | Konso | 36 | 13.96 | 43.51 | 29.55 | - | - |
| 18 | IRD | Guinea | Founi Ba | 36 | 14.70 | 43.68 | 28.98 | - | - |
| 19 | IRD | Guinea | Sira Rane | 38 | 14.15 | 41.75 | 27.60 | - | - |
| 25 | HNG | Guinea | Hotia | 34 | 14.64 | 43.48 | 28.84 | - | - |
| 26 | HNG | Guinea | Koine | 34 | 14.88 | 43.31 | 28.43 | - | - |
| 27 | HNG | Guinea | Konso | 37 | 14.46 | 41.67 | 27.21 | - | - |
| 28 | HNG | Guinea | Hotia | 35 | 17.99 | 45.03 | 27.04 | - | - |
| 29 | HNG | Guinea | Bambougou | 38 | 16.64 | 41.05 | 24.41 | - | - |
| 30 | HNG | Guinea | Rane | 38 | 16.90 | 40.77 | 23.87 | - | - |
| 31 | HNG | Guinea | Bode | 35 | 14.83 | 42.76 | 27.93 | -1.038 | -0.8 |
| 32 | HNG | Guinea | Rane | 38 | 16.33 | 40.08 | 23.75 | -1.0183 | - |
| 33 | HNG | Guinea | Kensere | 37 | 15.37 | 41.74 | 26.37 | - | - |
| 34 | HNG | Guinea | Nyougou | 35 | 11.98 | 43.45 | 31.47 | - | - |
| 35 | HNG | Guinea | Konso | 38 | 16.85 | 40.58 | 23.73 | - | -0.84 |
| 36 | HNG | Guinea | Bagala | 32 | 16.16 | 39.99 | 23.83 | - | - |
| | | | **Average** | 36.00 | 14.52 | 42.62 | 28.11 | -1.09 | -0.83 |

Figure S2 – Cardinal temperatures (optimum temperature, ceiling temperature, base temperature, and thermal window at t50) and base water potentials (Psib) at 30°C (t50) and 20°C (t15) for fonio seed accessions.

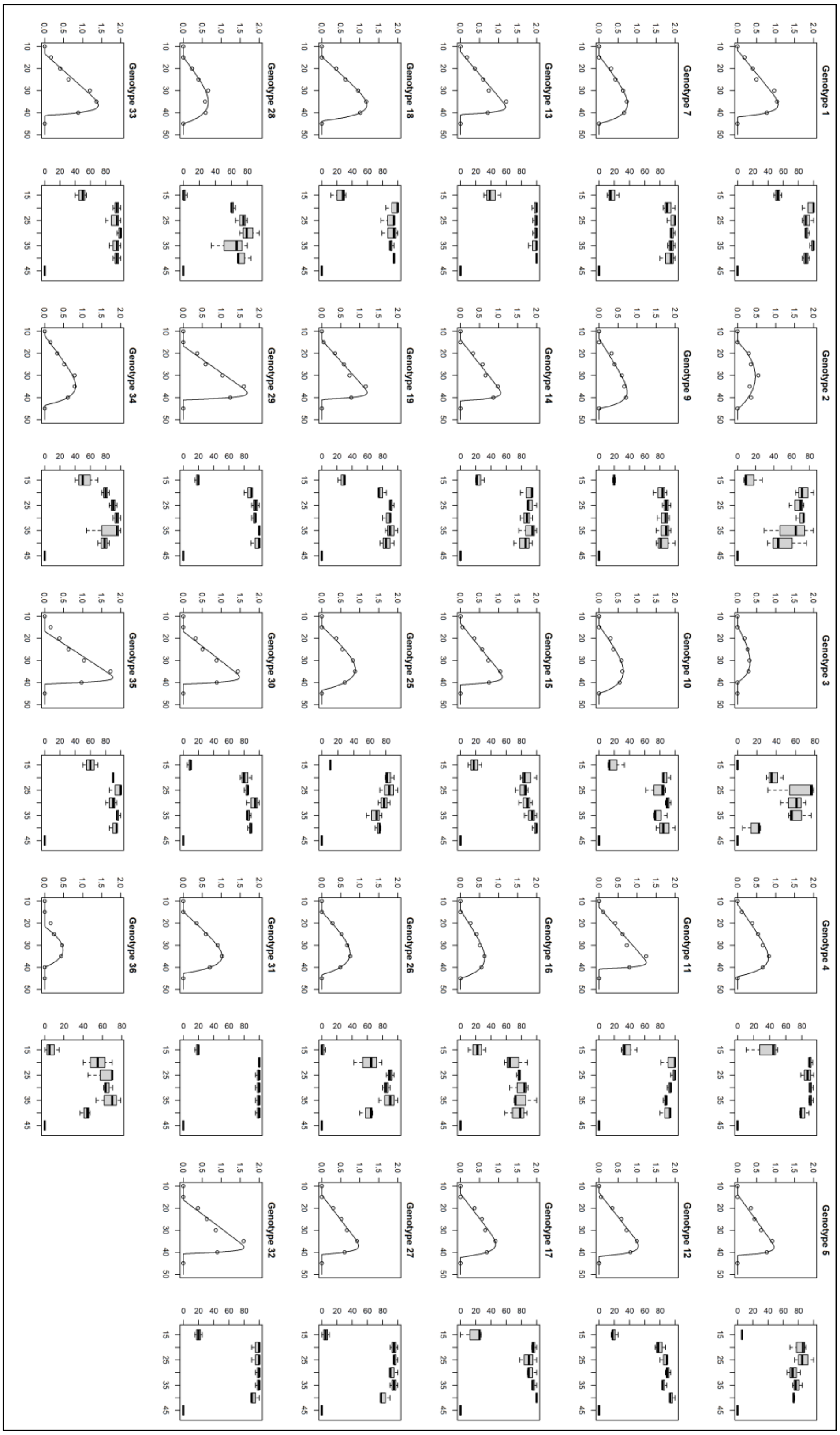


Figure S3 – Modelled germination rate curves and proportions (boxplots) for all accessions used in temperature experiments.

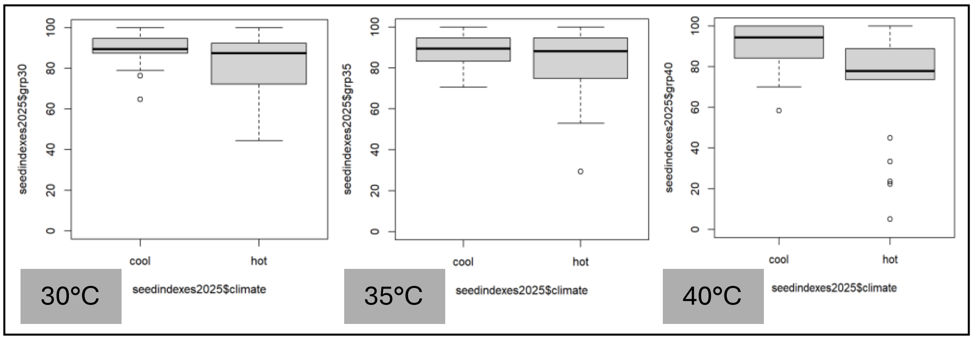


Figure S4 – Effect of seed accession origin climate (Guinea and Togo vs. Mali and Burkina Faso) on germination proportion at 30°C, 35°C, and 40°C.

| | Tw | Tc | Tb | To | Max Temp | Min Temp | Avg. Precip | Altitude | Latitude | grp(40) | grs(40) | gsp(40) | grp(35) | grs(35) | gsp(35) | grp(30) | grs(30) | gsp(30) |
|---|---|---|---|---|---|---|---|---|---|---|---|---|---|---|---|---|---|---|
| Tw | | | | | | | | | | | | | | | | | | |
| Tc | **2.74E-10** | | | | | | | | | | | | | | | | | |
| Tb | **2.05E-06** | 0.6278 | | | | | | | | | | | | | | | | |
| To | 0.6691 | 0.1592 | 0.3036 | | | | | | | | | | | | | | | |
| Max Temp | **0.008085** | 0.1558 | **0.02642** | **0.006238** | | | | | | | | | | | | | | |
| Min Temp | **0.03295** | 0.3728 | **0.03389** | **0.002782** | | | | | | | | | | | | | | |
| Avg. Precip | **0.004235** | 0.121 | **0.01815** | **0.001641** | | | | | | | | | | | | | | |
| Altitude | **0.006198** | 0.6266 | **0.0003** | **0.01612** | | | | | | | | | | | | | | |
| Latitude | **0.01636** | 0.9984 | 0.1126 | 0.408 | | | | | | | | | | | | | | |
| grp(40) | 0.05476 | 0.331 | 0.08824 | **6.93E-09** | **0.03933** | 0.1166 | 0.2601 | 0.2115 | 0.3722 | | | | | | | | | |
| grs(40) | **0.02414** | 0.2424 | 0.05059 | **1.75E-05** | 0.4925 | 0.8134 | 0.4355 | 0.5642 | 0.7084 | | | | | | | | | |
| gsp(40) | 0.5904 | 0.9433 | 0.4683 | 0.2842 | 0.4983 | 0.1853 | 0.3049 | 0.4249 | 0.2575 | | | | | | | | | |
| grp(35) | 0.4487 | 0.6985 | 0.1052 | **3.32E-06** | 0.3275 | 0.4242 | 0.667 | 0.2963 | 0.6521 | | | | | | | | | |
| grs(35) | 0.4426 | 0.7094 | 0.1057 | **2.96E-05** | 0.3906 | 0.7153 | 0.5387 | 0.204 | 0.5628 | | | | | | | | | |
| gsp(35) | 0.3871 | **0.04127** | 0.272 | **8.78E-07** | 0.09659 | 0.2982 | 0.2609 | 0.5095 | 0.67 | | | | | | | | | |
| grp(30) | **0.04793** | 0.334 | 0.07245 | **6.66E-05** | 0.296 | 0.4135 | 0.8201 | 0.8253 | 0.6137 | | | | | | | | | |
| grs(30) | 0.2705 | 0.9935 | 0.09282 | **0.000303** | 0.118 | 0.3186 | 0.8561 | 0.6079 | 0.44 | | | | | | | | | |
| gsp(30) | 0.574 | 0.07729 | 0.2138 | **7.46E-05** | 0.09402 | 0.6789 | 0.3559 | 0.4463 | 0.4897 | | | | | | | | | |
| Climate | 0.08215 | 0.3311 | 0.1518 | **0.0009284** | **<2.2e-16** | **2.52E-12** | **2.44E-07** | **7.57E-07** | **2.63E-12** | **5.23E-04** | 0.07981 | 0.4229 | 0.05428 | 0.09365 | **0.01641** | **0.00876** | **0.002714** | **0.03729** |
| Country | **0.01547** | 0.126 | 0.1643 | **2.09E-09** | **<2.2e-16** | **1.59E-13** | **9.25E-07** | **2.53E-10** | **<2.2e-16** | **9.80E-06** | **0.008964** | 0.8702 | **9.80E-06** | **0.008231** | **0.04088** | **0.00207** | **0.006768** | 0.1834 |

Figure S5 – Correlation of variables between seed germination proportions, cardinal temperatures, and climatic variables. Significant p values are in bold.

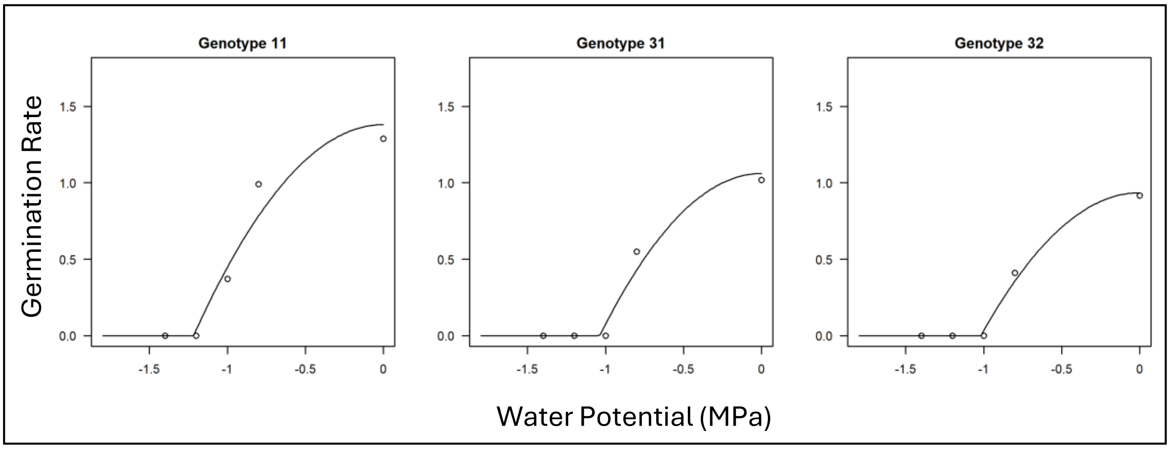


Figure S6 – Modelled germination rate of three fonio accessions under a gradient of water potential conditions, at threshold t50.

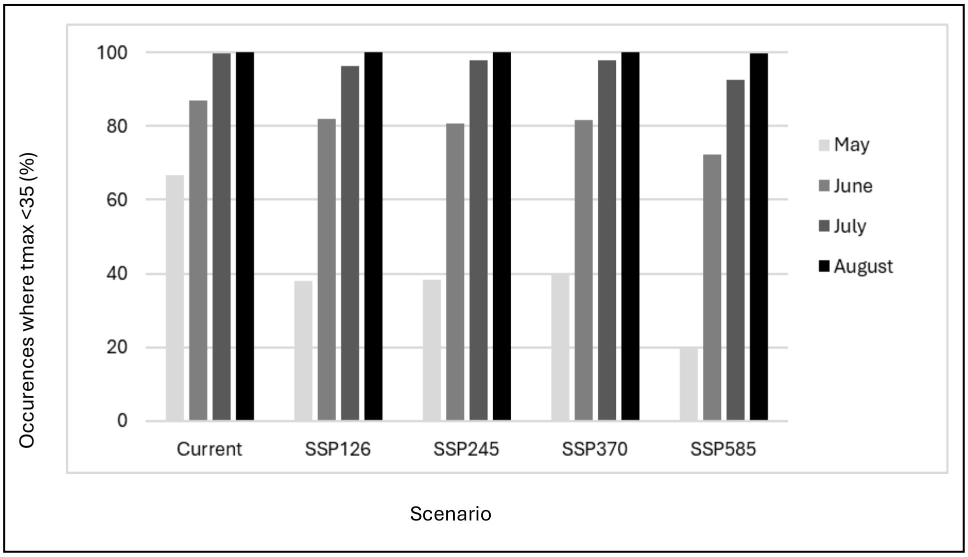


Figure S7 – Percentage of fonio occurrence points mapped to maximum temperatures above 35°C, in the current and under future climate SSP scenarios, grouped by month.

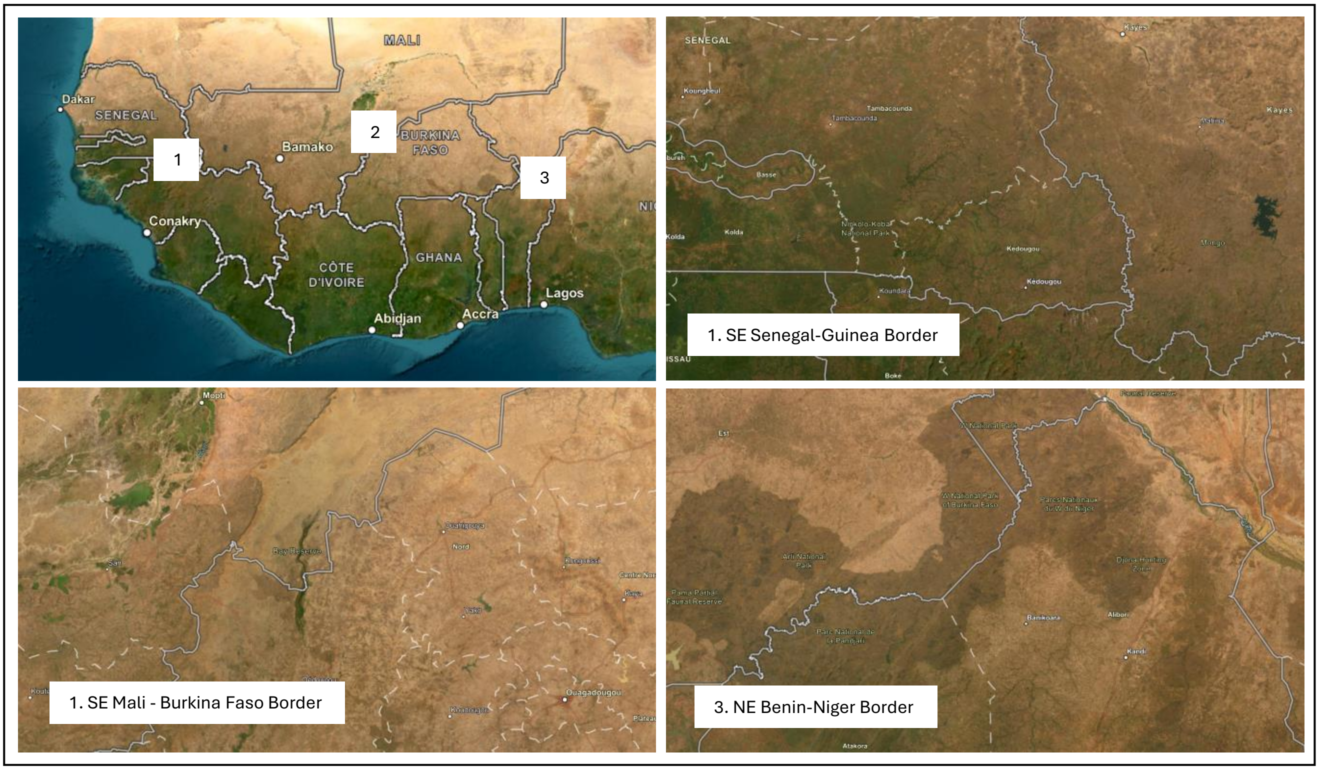


Figure S8 – Key areas predicted to experience significant reduction in suitable land for fonio cultivation.
